# Do it yourself: Creating 3D brain-surface models and custom-made brain matrices for guided sectioning using photogrammetry and three-dimensional printing technology

**DOI:** 10.1101/2025.01.27.635028

**Authors:** Sophie Holtz, Julio Hechavarria, Mirjam Knörnschild, Constance Scharff

## Abstract

Neuroethologists study many non-model animals. The development of techniques for precise and standardized histological brain analysis is key for understanding neural mechanisms across species. Here we present a novel cost-effective approach combining photogrammetry and three-dimensional printing technology to generate 3D brain surface models and produce species-specific brain matrices for precise and reproducible trimming, blocking, and sectioning of brain tissue. Using this strategy, we generated the first 3D brain surface model and brain matrix for Seba’s short-tailed fruit bat *Carollia perspicillata, CP*. We generated the 3D brain surface model with the open-source 3D graphics suite Blender and its inexpensive photogrammetry plug-in Snapmesh. High-quality photographs of two male *CP* brains were used for the 3D-reconstruction, resulting in a 3D surface model. Photogrammetry thus offers a viable, cheap alternative to techniques otherwise requiring access to costly 3D- or CT-scanners. The brain matrix was then modeled using Blender, and printed using a 3D printer. Our brain matrix facilitates and standardizes trimming or blocking of brains, providing consistent access to brain regions of *CP* in the sagittal and coronal planes. Using this workflow, brain surface models and brain matrices can be produced from brains without tissue loss. Our approach provides an affordable, and tissue-preserving alternative for the generation of 3D brain surface models and brain matrices and will benefit researchers globally, especially those with limited funding, no access to scanners or working with particularly valuable specimen.

## 1. Introduction

Neuroethologists study the brains of many different species, reflecting the broad spectrum of behaviours under investigation. The variation in brain size, shape, and anatomy of these non-model animals provides a challenge, as most methods, techniques, tools, and protocols used for standardized brain analysis are primarily designed for standard animal models. For instance, high-quality reference brain atlases exist for mouse (mouse.brain-map.org and atlas.brain-map.org), rats (Paxinos and Watson (2006), drosophila (Court et al. (2023); Schlegel et al. (2024) and zebra fish (Ullmann, Cowin, Kurniawan, and Collin (2010). Despite the growing interest in the development of standardized techniques and tools, such as reference brain atlases, for non-model animals, the required resources often exceed those of many research groups (Ron et al., 2024). Without standardized tools and techniques tissue analysis can vary, affecting reproducibility and reliability. It is therefore crucial to employ techniques for brain analysis that are precise and species-specific on the one hand, yet standardized on the other. Standardization of initial steps of histological brain analysis, such as tissue blocking or sampling specific brain regions, are especially critical. Utilizing brain matrices can stabilize the brain and guide trimming blades, significantly enhancing sectioning and sampling consistency, without requiring special neuroanatomical training (Defazio, Criado, Zantedeschi, & Scanziani, 2015; Huang, Chen, Fang, Chen, & Hung, 2019; Morawietz et al., 2004). Since matrix guided trimming approaches yield reproducible results (Defazio et al., 2015), they can also provide an accessible tool to share tissue samples with other researchers (Huang et al., 2019). Unfortunately, brain matrices are available only for comparatively few animals (Huang et al., 2019). Consequently, researchers working with non-model organisms often resort to freehand trimming, which can lead to sectioning and sampling inconsistencies, particularly for those with limited experience (Defazio et al., 2015; Huang et al., 2019; Morawietz et al., 2004).

To enhance reproducibility and reduce variability of sectioning for studies involving non-model organisms, one approach is to standardize the tissue embedding process. Currently, this approach relies on the use of specialized embedding chambers (Radtke-Schuller, Fenzl, Peremans, Schuller, & Firzlaff, 2020; Radtke-Schuller et al., 2016), where accurate brain positioning can be challenging. However, for standardized blocking, trimming, or sampling of specific brain regions without embedding, brain matrices are optimal. The generation of such a matrix requires two steps, creating a 3D model of the brain and subsequently creating the brain sectioning matrix using the model. This process generally requires access to CT or 3D scanners and 3D modelling software, and carries the risk of damaging valuable brain samples or significantly limiting downstream analysis options due to tissue handling in the model generation (Huang et al., 2019). These drawbacks are particularly relevant for resource limited researchers, be the resource money or the specimen availability. Addressing these challenges will thus also promote equity in science.

Here we describe an alternative workflow that neither relies on expensive scanning equipment for the generation of the 3D brain surface model, nor compromises tissue integrity during generation of the model or matrix. Instead it employs photogrammetry instead of 3D scanning, along with open-source or inexpensive software. Photogrammetry is a technique for deriving metric information about real-world objects by taking measurements from images, such as photographs or radar images, of those objects (Linder, 2013; Mikhail, Bethel, & McGlone, 2001). This method can also be used to reconstruct 3D shapes and is well established in geosciences, including archeology (Marín-Buzón, Pérez-Romero, López-Castro, Ben Jerbania, & Manzano-Agugliaro, 2021), cultural heritage documentation (Fritsch, Becker, & Rothermel, 2013; Hanan, Suwardhi, Nurhasanah, & Santa Bukit, 2015), as well as other areas of geological research (Eltner et al., 2016). Only recently has this technique become more accessible through various apps and software platforms, finding new applications in medical research as is provides a means for non-invasive 3D scanning at lower costs (Struck, Cordoni, Aliotta, Pérez-Pachón, & Gröning, 2019). In neuroscience, photogrammetry has been used to aid in head digitization for localization of scalp-mounted neuroimaging sensors (Clausner, Dalal, & Crespo-García, 2017; Mazzonetto, Castellaro, Cooper, & Brigadoi, 2022), to generate 3D surface models of autopsied human brains before subsequent analysis (Shintaku et al., 2019) and to aid in the collection of metric information about white matter tracts in the human brain (De Benedictis et al., 2018; Nocerino et al., 2017). However, photogrammetry’s reliance on features on the object’s surface makes it challenging to use on featureless objects that have smooth, texture-less, and glossy surfaces (Clausner et al., 2017). Consequently, it is not surprising that this technique has, to our knowledge, not been used to reconstruct 3D shapes of small vertebrate brains.

Here we demonstrate that photogrammetry can be used to generate 3D brain surface models of small vertebrates and that these models can be used to create species-specific matrices for guided brain sectioning, trimming or blocking. By combining photogrammetry for 3D brain surface reconstruction with 3D modelling and 3D printing technologies, using an open-source software and a cheap photogrammetry plugin, we have created the first 3D brain surface model as well as a species-specific brain sectioning matrix for Seba’s short-tailed fruit bat *CP*. Our approach provides an affordable and tissue-preserving technique to generate 3D brain surface models and brain matrices for small vertebrates, which could significantly benefit the neuroscience community globally.

## 2. Material and Methods

### 2.1 Animals

The brains of four adult male *CP* bats from the breeding colony of the Institute for Cell Biology and Neuroscience (Frankfurt University) were collected for a neurophysiology study approved by the Regierungspräsidium Darmstadt, Germany (permit number: FR/2007).

### 2.2 Perfusion and tissue processing

The animals were deeply anesthetized via an intraperitoneal injection of a mixture of Ketavet (active ingredient: 5 mg/mL ketamine hydrochloride; 7.5 mg/kg ketamine hydrochloride) and Rompun (active ingredient: 11 mg/mL xylazine hydrochloride; 16.5 mg/kg xylazine hydrochloride) and then euthanized with an overdose of sodium pentobarbital (400mg/1000g body weight; 160 mg/mL, Narcoren, Boehringer-Ingelheim, Ingelheim am Rhein, Germany), followed by transcardial perfusion with ice-cold 1x phosphate buffered solution pH 7.4 and 4% paraformaldehyde (PFA) in 0.1M phosphate buffer solution pH 7.4. For post-fixation, the bats’ heads were removed from the body, the skull slightly opened to expose the brain, and stored in 4% PFA in 0.1M PB pH 7.4 solution for 24h at 4°C. After post-fixation the brain was excised from the skull and the dura removed. The tissue was washed in 1x PBS and either stored in 1x PBS at 4°C C or subsequently cryoprotected by successive immersion in 15% and 30% sucrose solution in 1x PBS pH 7 before sectioning.

### 2.3 Creating the 3-dimensional brain surface model of CP brain using photogrammetry and subsequent generation of a brain matrix for sectioning

The workflow for creating a 3D brain surface model and generating the brain matrix for sectioning comprised three parts: I) generation of 3D surface brain surface model using photogrammetry, II) modeling of the brain sectioning matrix, and III) 3D printing of the brain sectioning matrix model (Figure 1).

**Figure 1:**
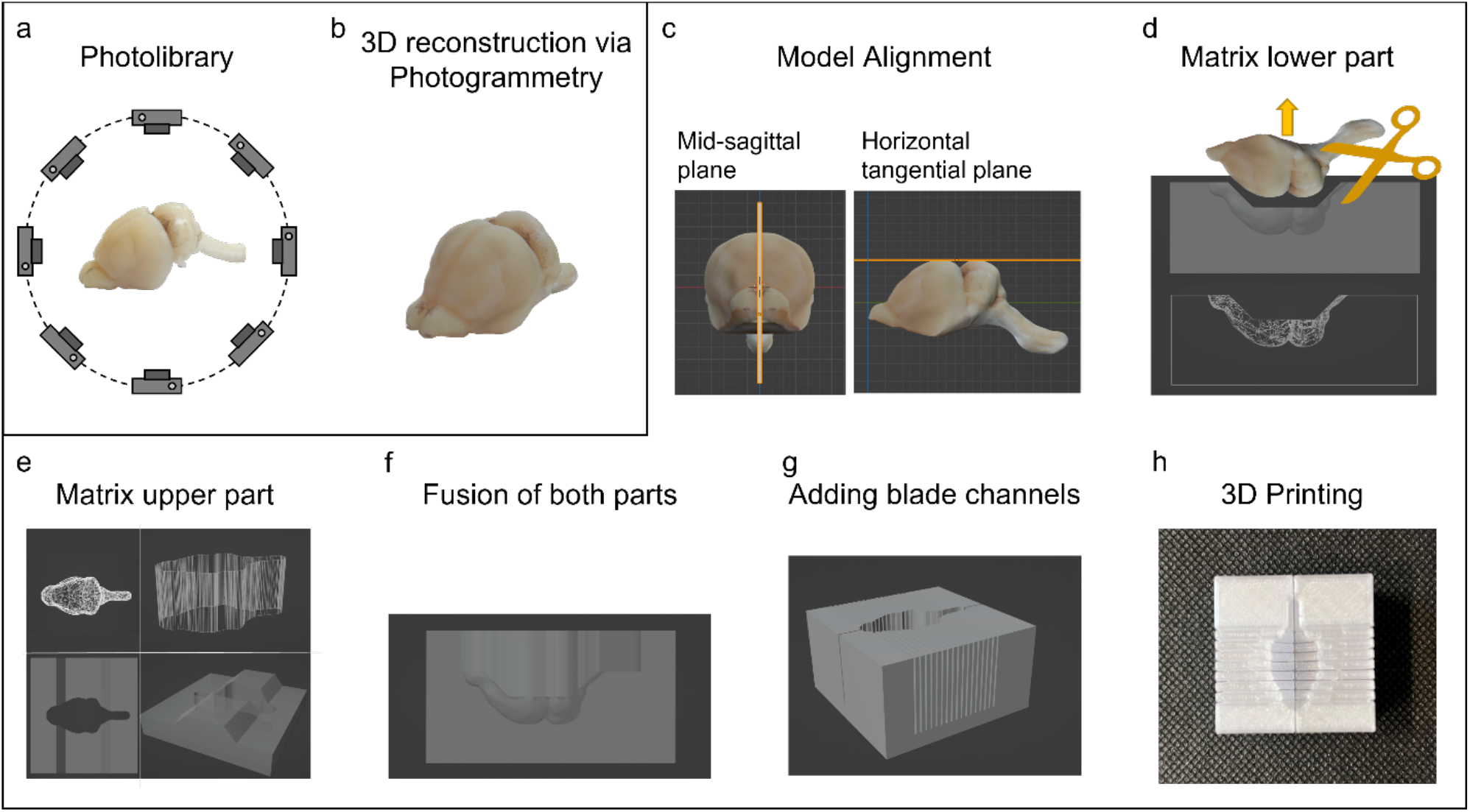
Workflow for the creation of the brain matrix for sectioning. A photo library containing pictures of the brain from all angles is compiled (a) and then used to reconstruct the 3D brain surface model using photogrammetry (b). The 3D brain surface model is subsequently aligned in space (c) and cut out from the lower part of the matrix cuboid (d). The maximum cross-sectional plane is determined by flat-top projection of the brain from dorsal view (e upper left), transformed into a 3D object (e upper right), and then cut out from the upper part of the matrix cuboid (e lower left and right). Upper and lower part of the cuboid matrix are fused into one brain matrix (f) and blade channels that guide blades through the brain are added (g). Finally, the 3D model of the brain sectioning matrix is printed using a 3D printer (h). For detailed workflow of the matrix modeling see Figure 2.

#### I) Generating a 3D brain surface model

Creating a photo library of the brains

Photogrammetric reconstruction relies on spatial cues on the surface of the object and the area around the object. The quality of a 3D surface model generated using photogrammetry depends on the quality of the photo library used for reconstruction. Polarizing filters and consistent lighting conditions help to minimize reflections, highlights, and shadows that result from the shininess of the brain surface. Photographs should be taken from enough different viewpoints covering all aspects of interest of the object while keeping sufficient overlap between neighboring picture pairs. The optimal changes in angle and the rotation steps between the photographs depend on the object’s geometry (De Paolis, De Luca, Gatto, D’Errico, & Paladini, 2020). The photographs should be taken with a deep depth of field to keep as much of the brain as possible in focus to prevent that only parts of images are sharp enough to be used in the 3D reconstruction process (De Paolis et al., 2020). Alternatively, pictures with different focal distances can be combined using focus stacking techniques, resulting in a single picture with a higher depth of field, that can be used for photogrammetry (Clini, Frapiccini, Mengoni, Nespeca, & Ruggeri, 2016; De Paolis et al., 2020; Kontogianni, Chliverou, Koutsoudis, Pavlidis, & Georgopoulos, 2017).

In our study, the brains of two adult male *CP* bats were placed ventral side down on a white piece of paper. We blotted excess moisture from the brains to reduce shininess, and placed markings on the piece of paper as additional spatial cues. Using a DXLR camera (Canon EOS 550D) with a 55 mm lens (Canon EF-S 18-55 mm F/3.5-5.6 IS STM) we took 120-140 pictures per brain within 5-7 minutes to avoid tissue desiccation. Photos were taken from all sides except the ventral side, using different angles by moving the camera around the brains (Figure 1A).

##### 3D Reconstruction using photogrammetry

A total of 260 photos from both *CP* brains were employed for the 3D reconstruction to generate the 3D surface model (Figure 1B and Figure 3). We used the open-source graphics suite Blender and its commercially available, inexpensive, photogrammetry plug-in SnapMesh (settings: details = raw, feature sensitivity = high, object masking = true).

#### II) Modeling of the Brain Matrix for sectioning

The brain matrix was modeled in two halves using Blender (Figure 1c-g); a lower half and an upper half that were later fused together.

The 3D brain surface model was oriented in space with its midsagittal plane aligned orthogonal to the horizon, its horizontal tangential plane defined by the most dorsal elevation of cerebellum and cerebrum, parallel to the horizon (Figure 1c), and its dorsal side pointing down.

Next, a cuboid was aligned in space around the 3D brain surface model with its ventral and dorsal surface parallel to the horizon (Figure 2a). The cuboid was big enough to fit the entire 3D brain surface model while keeping a minimum distance of 4 mm millimeter between the brain model and all surfaces of the cuboid, which translates into the wall-thickness of the matrix around the brain chamber. This was to ensure stability for the brain matrix even after adding blade channels, which run through the whole brain chamber into the matrix body, and generate a final minimal wall-thickness of 1 mm for the brain sectioning matrix.

**Figure 2:**
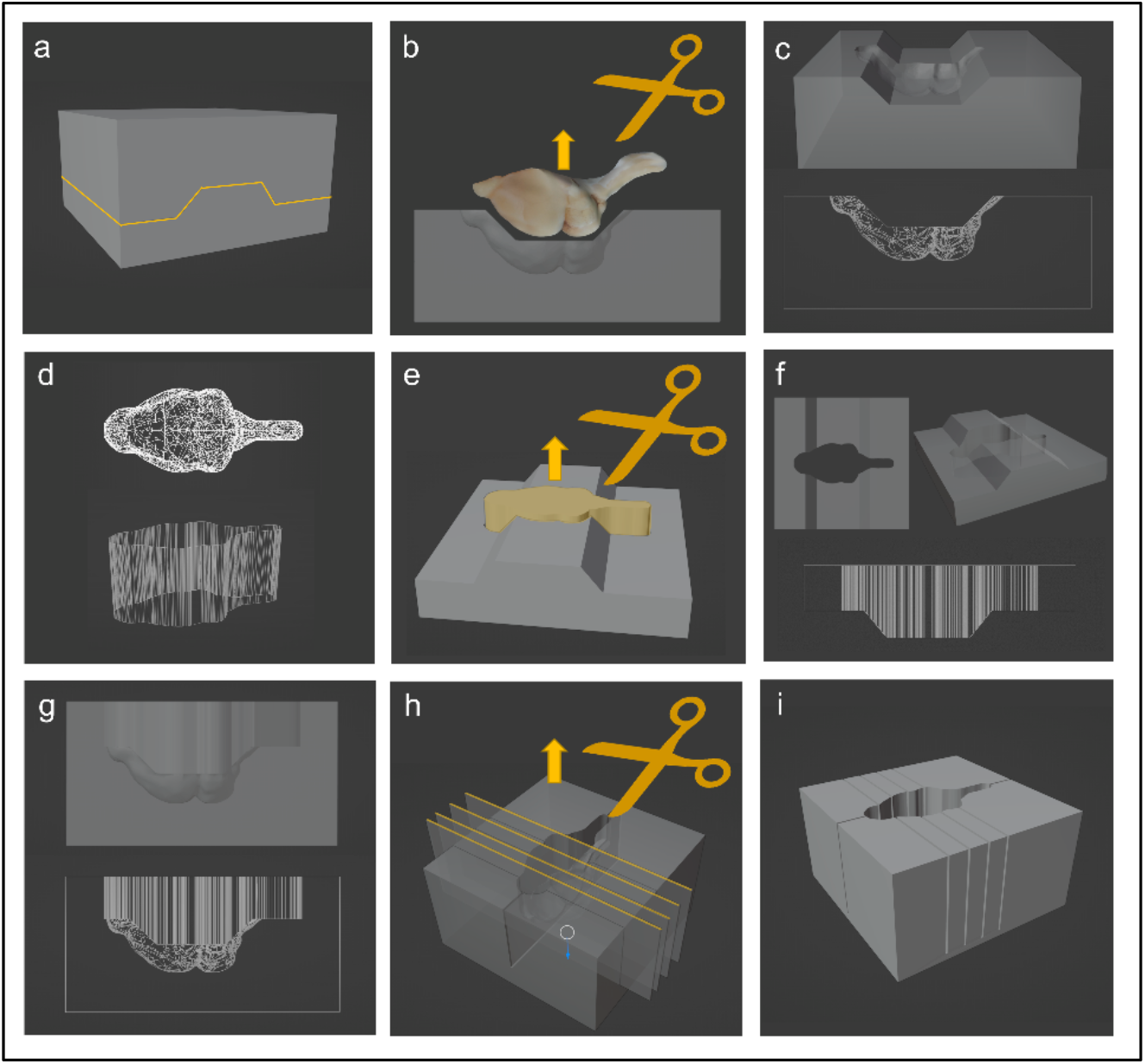
Detailed workflow for modelling of the brain sectioning matrix. A cuboid, aligned with the 3D brain surface model, is cut into two halves with the cutting line (yellow line) following the maximum cross-sectional points from a dorsal view (a). The 3D brain surface model is cut out from the lower part (b) to generate the lower part of the brain sectioning matrix (3D and wireframe representation, c). A 2D flat top projection of the brain from dorsal view (wire frame representation, d top) is transformed into a 3D object (wire frame representation, d bottom) and then cut out from the upper cuboid part (e) to generate the upper part of the brain sectioning matrix (3D and wireframe representation, f). Upper and lower part are fused together to create the whole brain sectioning matrix (3D and wire frame representation, g). Flat cuboids with measures of blades intended to be used for brain blocking or trimming are cut out from the brain matrix at desired locations (h) to generate the final brain sectioning matrix (i).

**Figure 3:**
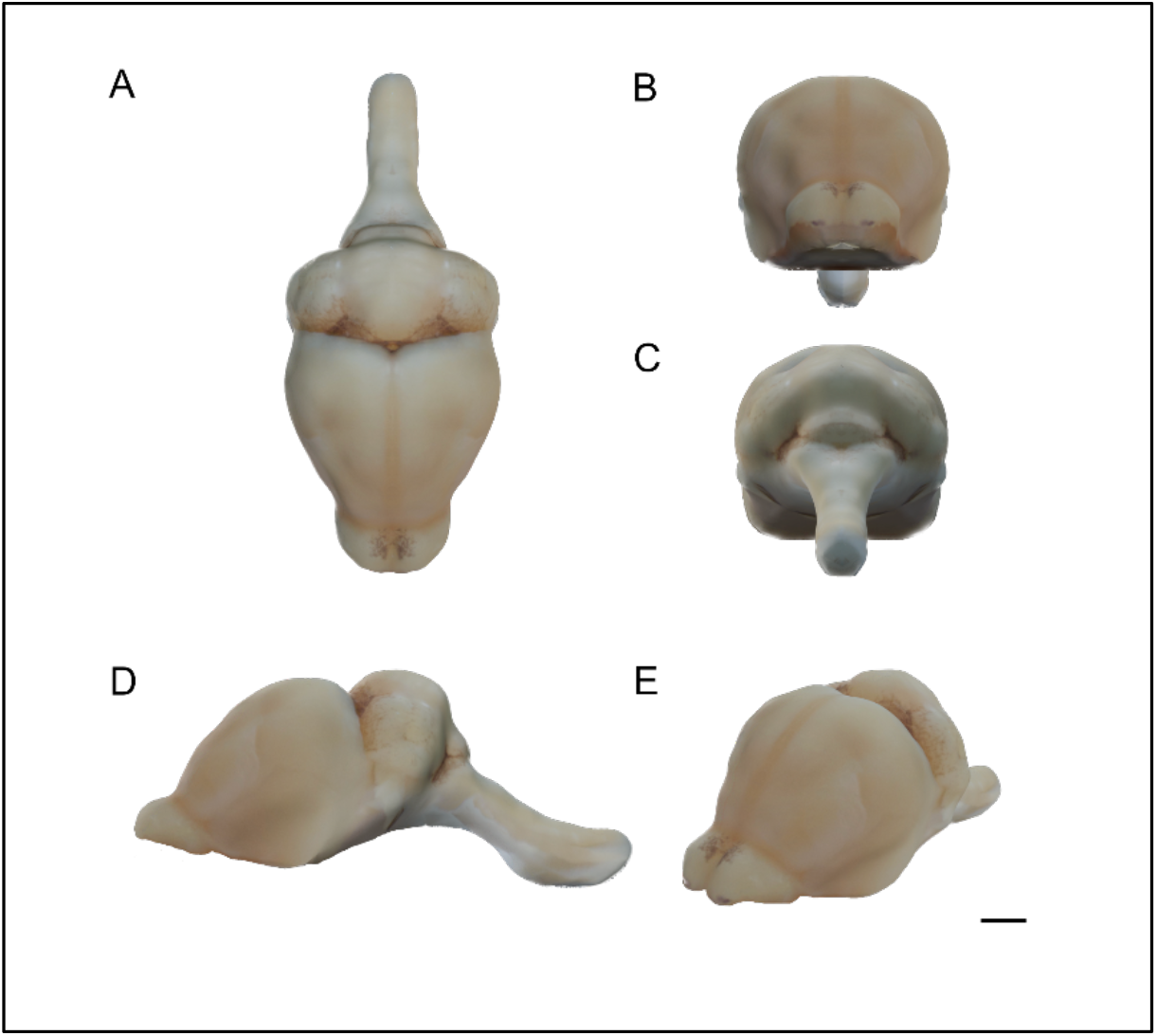
**3D reconstruction of the brain of CP** from dorsal/top-(A), anterior-(B), posterior-(C), and lateral view (D) and in semi profile(E). (Photogrammetry software settings: Details = raw, Feature sensitivity = high, Object masking = true). Size bar = 2 mm.

To be able to place a brain in the brain chamber of the sectioning matrix and remove it, the maximum cross-sectional plane of the brain (i.e. the widest points of the brain determined by a 2D flat top projection of the brain model in dorsal view, Figure 2d left, 1e top left) was determined and the cuboid was horizontally cut in two halves creating a lower and an upper half, with the cutting line roughly following points marking the maximum cross-sectional plane along the brain in a lateral view (Figure 2a). Afterwards, the 3D brain surface model was cut out from the bottom half using Blender’s Boolean difference modifier (Figure 1d and 2b) resulting in the lower half of the brain matrix (Figure 2c).

To create the top half of the sectioning matrix (Figure 1e and 2d-f), the outline of the 2D flat-top projection of the 3D brain surface model (Figure 2d left) was used to create a 3D object of the brain’s maximum cross-sectional outline (Figure 2d right). The resulting 3D object was then cut out from the top half using blender’s Boolean difference modifier (Figure 2e), resulting in the top half of the brain sectioning matrix (Figure 1e, 2f). Because 2D and 3D object were created from the already aligned 3D brain surface model, they were already correctly aligned with the cuboid and did not have to be repositioned during this process. Next, lower and upper half were fused together using blender’s Boolean union modifier (Figure 1 F and 2g). To be able to place a brain inside the matrix and remove it without damaging the tissue, the size of the matrix model was increased by 1 mm in all dimensions. Size adjustments for the matrix should be made before adding blade channels, to not alter dimensions of the channels, only the matrix.

Finally, blade channels that later guide the trimming blade through the brain were added to the matrix model (Figure 1g and 2h). Cuboids with a length and height greater than the matrix dimensions, and a width of 0.2 mm were cut out from the brain matrix (Figure 2h), leaving a minimum distance of 1 mm to the ventral surface of the matrix to ensure stability. The width of the blade channels was chosen to fit the thickness of the blades used to cut brains. We added one mid-sagittal blade channel to provide the means to separate both hemispheres and create a sagittal sectioning plane, as well as four frontal blade channels (Figures 2h - I, and 5). Number, spacing, as well as orientation and angle of the blade channels can be customized to accommodate specific research questions, such as to sample specific brain regions.

#### III) 3D printing of the brain matrix model

For 3D printing, the 3D brain matrix model was printed using a Bambu Lab X1C 3D printer and PETG filament (Figure 1h, 5).

### 2.4 Assessment of the model’s quality

To determine variability in brain sizes and shapes across the brains used for 3D reconstruction and to compare them with the 3D surface model for quality assessment of the model, measurements were taken at easily identifiable landmarks along the lateral-mesial axis and at various points from dorsal and lateral views (Figure 4B, Table S1). Brain width was measured at six landmarks: the widest point of the olfactory bulb (1), the coronal suture running through Bregma (2), along the line running through Lambda (3), at the Lambda suture (4), at the cerebellum (5), and at the medulla oblongata where the cerebellum ends (6). Distances between landmarks were determined from both, dorsal and lateral views. In dorsal view, measurements were taken between the most frontal tip of the olfactory bulb (A), the most frontal point of the cerebral cortex (B), Lambda (C) and at the medulla oblongata where the cerebellum ends (D). From lateral view distances were measured between these landmarks (A-C, A-D, and C-D) and an additional point E, the most ventral point of the cerebral cortex (A-E, C-E, and D-E). Measurements were taken using Blender’s measurement tool for the 3D model, and photographs of the brains on millimeter paper for the brains.

**Figure 4:**
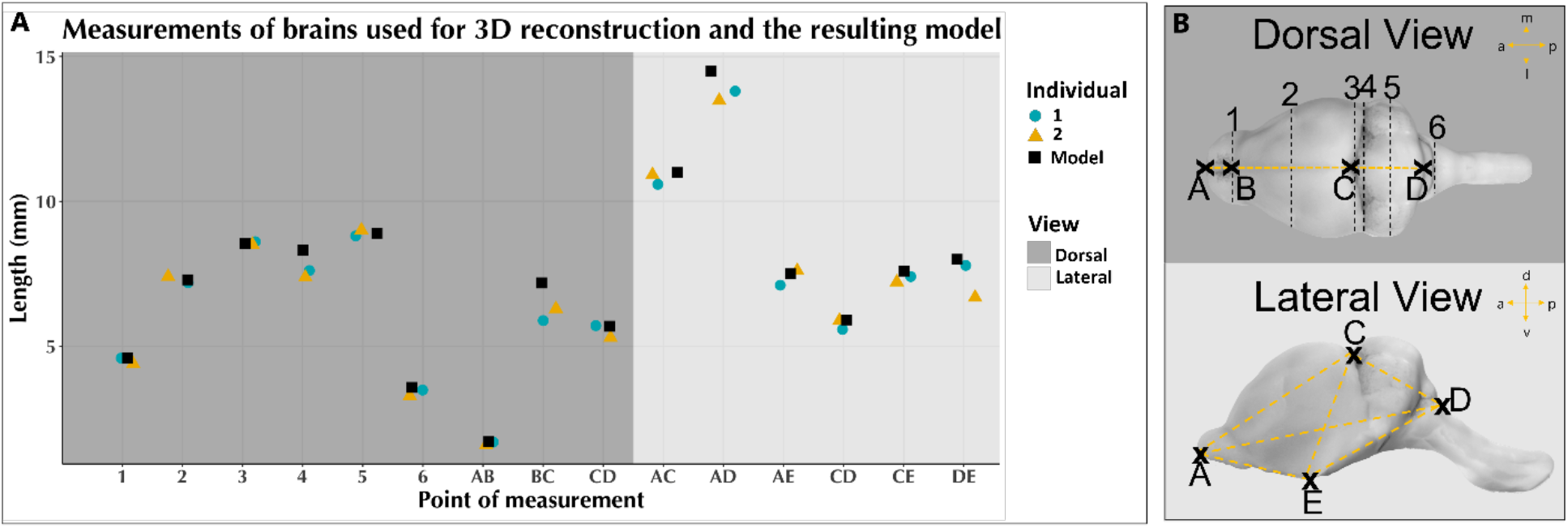
**Comparison of CP brains and 3D brain surface model.** Measurements across individual CP brains (yellow triangle and blue circle) used for reconstruction and the 3D brain surface model (black square) were taken from dorsal (dark grey shading) and lateral (light grey shading) view (A). Locations of points of measurement are shown on the 3D brain surface model (B). The brains width was measured at six different locations (1-6): 1 = Olfactory bulb, 2 = coronal suture running through Bregma, 3 = line running through Lambda, 4 = Lambda suture, 5 = Cerebellum, 6 = Medulla oblongata where the cerebellum ends. Distances between landmarks A (Tip of the olfactory bulb, most frontal point of the brain), B (most frontal point of the cerebral cortex), C (Lambda), D (endpoint of the cerebellum), and E (most ventral point of the cerebral cortex) on the brain. Scalebar = 2 mm 386

### 2.5 Matrix-guided blocking and successive Microtome sectioning and Nissl staining

We evaluated the quality of the sectioning matrix using brains of two additional adult *CP* males. The brains were processed as described in Methods, placed dorsal side down into the brain sectioning matrix and blocked by a frontal cut using blade channel number 3 (Figure 5 yellow dotted line) with a razor blade (Gillette Super Silver, width: 2 mm). Each brain block was then sectioned at 40 μm thickness using a freezing sliding microtome (Leica SM 2000 R) and collected in 24-well plates, so that each section had a unique number. Sections were stored in 30% ethylene glycol and 30% sucrose in 0.2M PB at -20°C.

**Figure 5:**
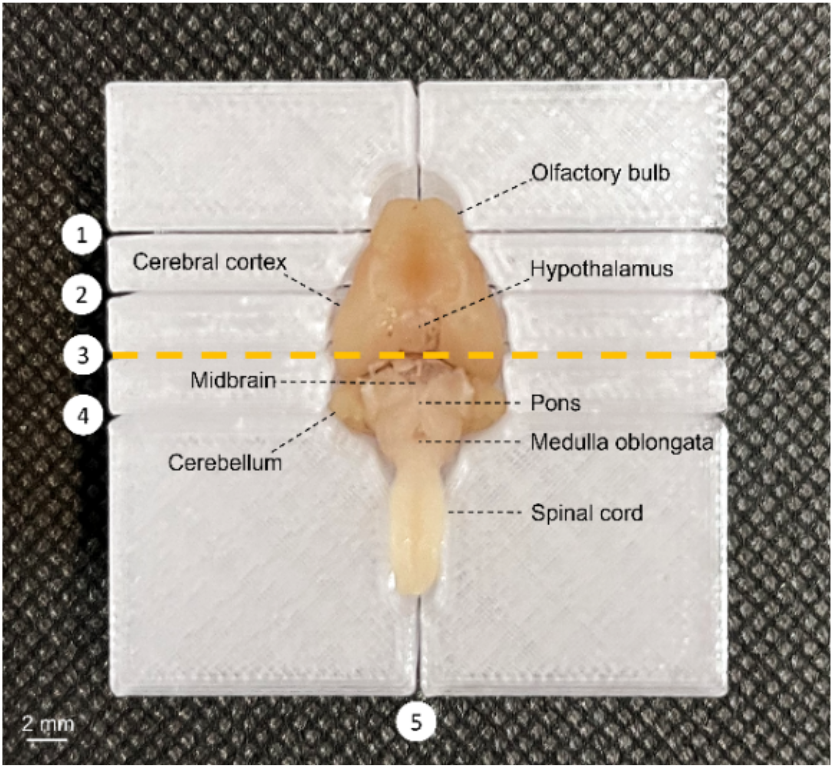
**Brain of an adult male CP in the brain sectioning matrix** with annotated anatomical landmarks. The matrix facilitates blocking, trimming or sampling in sagittal (blade channel 5) and coronal (blade channel 1-4) orientation. Coronal blade channel 3 (yellow dashed line) was used to generate brain blocks for subsequent microtoming in this study. Size bar = 2mm.

To assess the standardization achieved by using the matrix to block brains, brain sections at comparable anatomical locations, e.g. with the same section number, were selected for Nissl staining. Because the olfactory bulbs and the spinal cord of both brains were preserved up to different levels after dissection, section numbers for anterior and posterior brain blocks were counted from the start of the cerebral cortex and the cerebellum, respectively.

The selected sections were mounted onto chrome alum gelatin-coated glass slides, air-dried overnight, and stained with 0.13% Cresyl violet. Sections were delipidized using Xylene (5min, twice) and 100% Ethanol (2min), followed by sequential rehydration using decreasing ethanol concentrations (100% 2min; 2min 95%, twice; 2min 70%; 2min 50%; distilled water). Subsequently the sections were stained in 0,13% Cresyl violet solution for 7 minutes. Afterwards, the sections were washed in distilled water, moved through the ascending EtOH row and the stain differentiated with dips in 95% ethanol with 1% acetic acid followed by dehydration in 100% ethanol, and cleared using Xylene (2min, three times). Directly after the last Xylene wash, the slides were coverslipped using Omnimount medium (17997-01 Electron Microscopy Sciences) and air dried overnight. The sections were photographed under a Leica microscope (MDG30) equipped with a Leica Z16 APO lens and a Leica DFC 420 C camera using Leica Application Suite (V 4.8). For annotation of the sections, published brain atlases for the bat species *CP* (Scalia, Rasweiler, Scalia, Orman, & Stewart, 2013), *Phyllostomus discolor* (Radtke-Schuller et al., 2020), *Rousettus aegyptiacus* (Eilam-Altstadter, Las, Witter, & Ulanovsky, 2021), and the Allen mouse brain atlas (mouse.brain-map.org and atlas.brain-map.org) were used as references.

## 3. Results

### Creation mof a 3-dimensional brain surface model and subsequent generation of a species-specific brain matrix for sectioning of CP brains

By using photogrammetry combined with successive 3D-modeling and printing, we successfully generated a 3D brain surface model for *CP* (Figure 3) and produced the first bat-specific brain matrix for precise and reproducible blocking and sectioning (Figure 5). This novel, cost-effective workflow proved to be accessible, easy to learn and did not harm the tissue of the brains used for the photo library, making it a viable alternative to existing protocols for generating 3D brain-surface models and brain sectioning matrices for other small mammalian study species.

A photo library of 260 pictures from two brains was sufficient to generate the 3D brain surface model (Figure 3). Since the photos were taken with the brains positioned on their ventral side, the reconstruction was successful for all aspects except the ventral part, which was not required for our goal to generate of a sectioning matrix for this species. However, additional photographs could be included in the reconstruction process to also cover the ventral surface of the brain.

### Quality assessment of 3D brain surface model and brain sectioning matrix

To evaluate the variability in brain shape and size, as well as the quality of the 3D brain surface model, measurements across both brains and the model were taken (Table S1, Figure 4). All landmarks used for these measurements could be determined on both the brains and the model. The brains and the 3D reconstruction compared well to each other, with only a few measures showing more variability. On average, the two brains differed by 0.31 mm (std 0.24) across all measurements. The biggest discrepancy between the brains was the distance between D and E from lateral view, measuring 7.8 and 6. 7 mm for the two individuals. Most measures of the model fell between (measures 2, 3, 5, AE) or matched those of the brains (1, AB, CD lateral and dorsal view), except for the measures 4, BC, and AD, where the model exceeded the larger of the two brains by 0.7 mm to 0.9 mm.

Regardless of these small deviations, the 3D surface model of *CP’s* brain was sufficiently accurate for creating a species-specific brain sectioning matrix (Figure 5). The sectioning matrix facilitates precise blocking and trimming of *CP* brains in both, frontal and sagittal orientation. While the sagittal blade channel running along the midline of the brain (channel 5) separates both hemispheres, the frontal blade channels (1-4) can be used to generate frontal sectioning planes, to hemi-sect the brain at different levels, or to sample specific parts of the brain. The position of blade channels can be customized easily to cater to the researchers needs, for example to border specific brain regions of interest for matrix guided sampling.

To further validate the 3D brain surface model and the sectioning matrix, brains from two additional adult *CP* males were blocked using blade channel 3 of the sectioning matrix prior to sectioning with a freezing sliding microtome (Figure 5 yellow dashed line) and Nissl staining (Figure 6). The brains could easily be placed in and removed from the matrix and remained stable during the blocking procedure. The blade channels maintained stability, accommodating the 0.2mm thick razor blade accurately. The brains of individuals A and B differed in overall shape, which can be observed for all selected sections (Figure 6). Nevertheless, the brain sectioning matrix did support both of them equally well during blocking and created brain blocks of equal quality.

**Figure 6:**
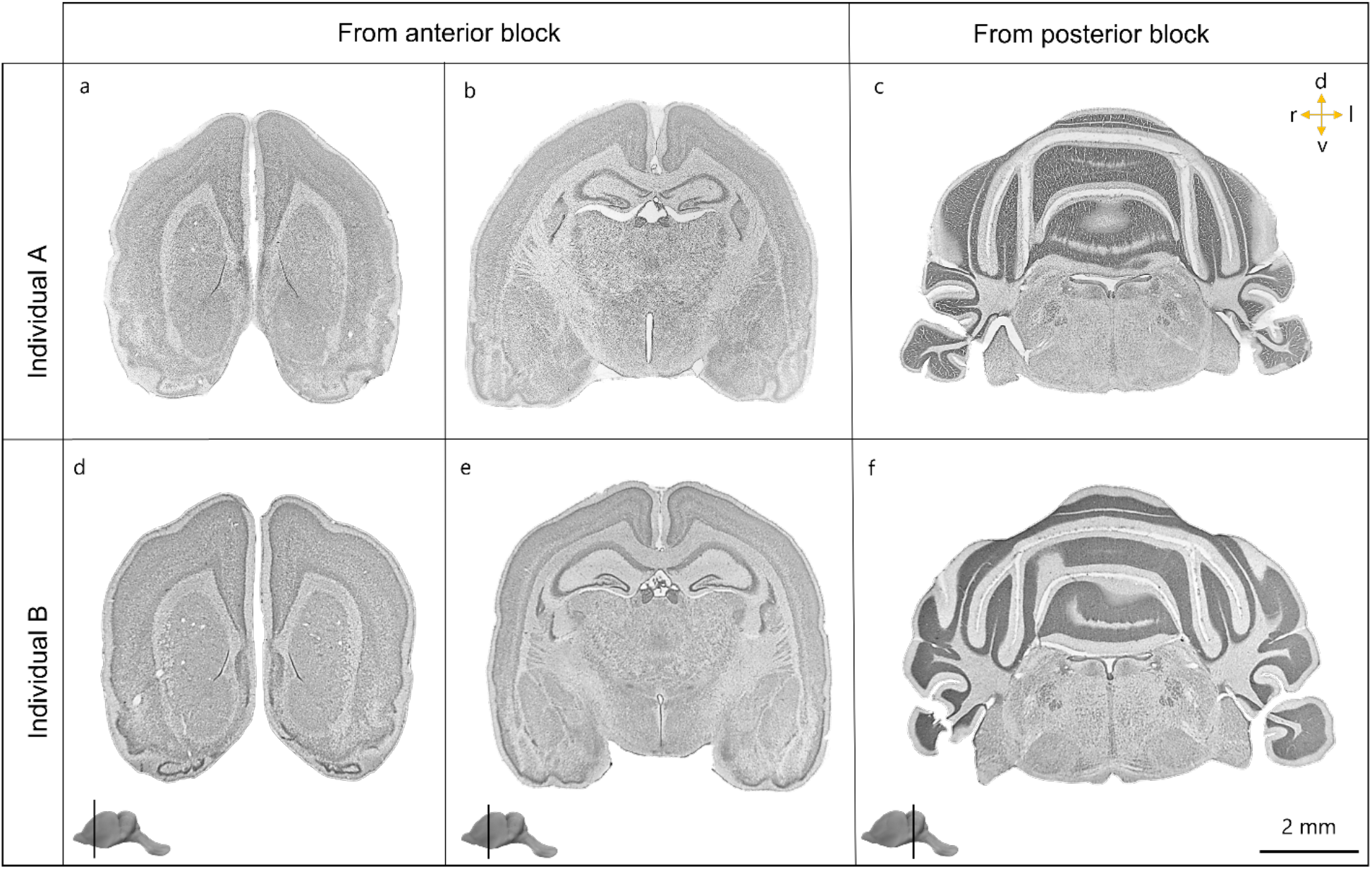
**Nissl-stained brain sections of two adult CP males** from anterior (a-b, d-e) and posterior (c, f) brain blocks generated by following the matrix guided blocking approach using the species-specific brain matrix followed by freezing microtoming. The brain blocks were generated using blade channel 3 of the sectioning matrix. Locations of the sections are shown. Size bar = 2 mm. For detailed annotations of the depicted brain sections see Supplementary Figures S1 to S6.

The stained sections from both, the anterior (Figure 6a-b and d-e) and posterior block (Figure 6c and f) showed high degrees of interhemispheric symmetry in both individuals, confirming correct alignment of the brain model in the matrix modeling process, resulting in accurate orientation of brains during sectioning. This also speaks to the quality of the 3D brain surface model. A slight left right asymmetry in individual B can be observed, which is most pronounced in the section of the posterior block at the level of the cerebellum.

The analysis of corresponding sections from both individuals revealed that corresponding section numbers generally showed the same anatomical regions, though individual B exhibited slightly more posterior aspects of these regions compared to individual A.

## 4. Discussion

In this study we demonstrated that the combination of photogrammetry, 3D modelling and -printing can be a viable and cost-effective alternative for the generation of 3D brain surface models and brain sectioning matrices for small vertebrates such as Seba’s short-tailed fruit bat *CP*. Our approach offers a promising alternative to traditional methods that use expensive equipment and software, likely benefiting researchers with limited resources.

To our knowledge, employing photogrammetry to generate 3D brain surface models has to date only been used on autopsied human brains (Shintaku et al., 2019). Here, we successfully applied this technique to small mammalian brains, although certain limitations need to be addressed. Photogrammetric 3D reconstruction relies on identifying reference points on the object’s surface that are shared between the photographs, but appear in different positions in each image on them (Shintaku et al., 2019). In the example of human brains, the brains’ surface structure provides numerous reference points. The often simpler and smoother brain surfaces of small vertebrates present far fewer of such features, particularly in perfused samples. Adding external reference points, such as markings around the object, or utilizing internal features like blood vessel patterns, can facilitate the 3D reconstruction process. In our case, rests of coagulated blood under the dura at different locations very likely served as additional reference points, enhanced reconstruction accuracy (Figure 3). If unperfused, immersion-fixed brains are available, potentially with the dura mater still attached this could improve the accuracy of the brain model further.

Our primary objective was to generate a brain sectioning matrix for this species, rendering a full reconstruction of the brain’s ventral aspect unnecessary. However, generating a complete 3D model can in principle be achieved by placing the brain on a transparent surface during imaging (as shown by Shintaku *et al*., 2019) or by combining separate models created from different perspectives. In our assessment of the 3D surface model’s quality, we observed some measurement deviations between the model and the actual brains, particularly for measures 4, BC, and AD. These discrepancies likely resulted from slight variation in the brain positioning and camera angles during imaging. These discrepancies could become particularly relevant when comparing measurements taken from photographs with those taken in Blender, where exact viewpoints may not perfectly align with those in the pictures. Such deviations also highlight the importance of producing brain matrices of different sizes to accommodate for size discrepancies between individuals both within and across age groups, as suggested by Huang et al (2019). While the deviations did not affect the quality of our brain sectioning matrix, it is important to consider them for other potential applications, like aiding in alignment of brain sections or scans within a common reference space. Given that the required precision of a model is highly dependent on its intended use, future research should further evaluate the accuracy of photogrammetric reconstructions against other 3D scanning methods, such as μCT, especially for those cases requiring high precision.

Photogrammetry is an evolving technique, with continuous advancements in soft- and hardware increasing precision and accessibility. The growing availability and affordability of photogrammetry tools, ranging from plug-ins and standalone software to smartphone apps, could make the routine reconstruction of individual brains feasible, adding valuable spatial data to downstream analyses, making photogrammetry a promising approach for future neuroanatomical research.

While the idea of using 3D printing to create custom sectioning matrices is not new, our approach leverages open-source and easy-to-learn software, such as Blender, to make the modeling process more accessible, customizable, and less resource intense. This enables researchers to easily adjust matrix dimensions and blade channel orientations to suit specific experimental needs and research questions. We produced our sectioning matrix using a cost-effective material comparable to stereolithography resin PETG filament, however its limited heat resistance constrains sterilization. Nevertheless, 3D matrix models can be printed in stainless steel if required.

Standardizing methods to ensure reliable and consistent results are essential for improving research efficiency and minimizing the number of animals required. Implementing a tissue-loss-free approach for generating sectioning matrices further supports this goal by reducing tissue waste, thereby decreasing the need for additional specimens.

Overall, our workflow offers a simplified, cost-effective, and tissue-loss free alternative for the generation of 3D brain surface models as well as matrices for guided brain sectioning to existing protocols that use costly 3D or CT scanners, expensive modeling software, and may result in tissue loss. Thus, our workflow could benefit researchers globally, especially those with limited financial resources or studying particularly valuable specimen.

## Author contributions

SH conceived, designed and performed all parts of the presented work, SH and CS wrote the manuscript, JH provided the animals, all authors consulted on various aspects of the work and all edited the paper. MK and JH provided feedback on the writing.

## Acknowledgements

We thank Tobias Hartwig for 3D printing and printing support, the members of Scharff lab, Knörnschild lab and Max Holtz for critical comments and reading. This project was supported by BMBF and Max Planck Society.

## Notes

### Competing Interest Statement

The authors have declared no competing interest.

