## Supplementary Material for "Do it yourself: Creating 3D brain-surface models and custom-made brain matrices for guided sectioning using photogrammetry and three-dimensional printing technology"

***Table S1: Measurements across the 3D brain surface model and the two individual CP brains used for the generation of the 3D brain surface model (mm).*** *Measures were taken from a dorsal view (dark grey shading) and a lateral view (light grey shading). The width of the brain was determined at locations 1-6: 1 = Olfactory bulb, 2 = coronal suture running through Bregma, 3 = line running through Lambda, 4 = Lambda suture, 5 = Cerebellum, 6 = Medulla oblongata where the cerebellum ends. Distances were measured between the landmarks A (Tip of the olfactory bulb, most frontal point of the brain), B (most frontal point of the cerebral cortex), C (Lambda), D (endpoint of the cerebellum), and E (most ventral point of the cerebral cortex). Measures taken from lateral view relate to a combination of length and height of the brains.*

| | Width at locations (mm) | | | | | | Distances between landmarks (mm) | | | | | | | | |
|---|---|---|---|---|---|---|---|---|---|---|---|---|---|---|---|
| | 1 | 2 | 3 | 4 | 5 | 6 | AB | BC | CD | AC | AD | AE | CD | CE | DE |
| **Individual 1** | 4,6 | 7,2 | 8,6 | 7,6 | 8,8 | 3,5 | 1,7 | 5,9 | 5,7 | 10,6 | 13,8 | 7,1 | 5,6 | 7,4 | 7,8 |
| **Individual 2** | 4,4 | 7,4 | 8,5 | 7,4 | 9 | 3,3 | 1,6 | 6,3 | 5,3 | 10,9 | 13,5 | 7,6 | 5,9 | 7,2 | 6,7 |
| **3D Brain Surface Model** | 4,6 | 7,3 | 8,55 | 8,3 | 8,9 | 3,6 | 1,7 | 7,2 | 5,7 | 11 | 14,5 | 7,5 | 5,9 | 7,6 | 8 |

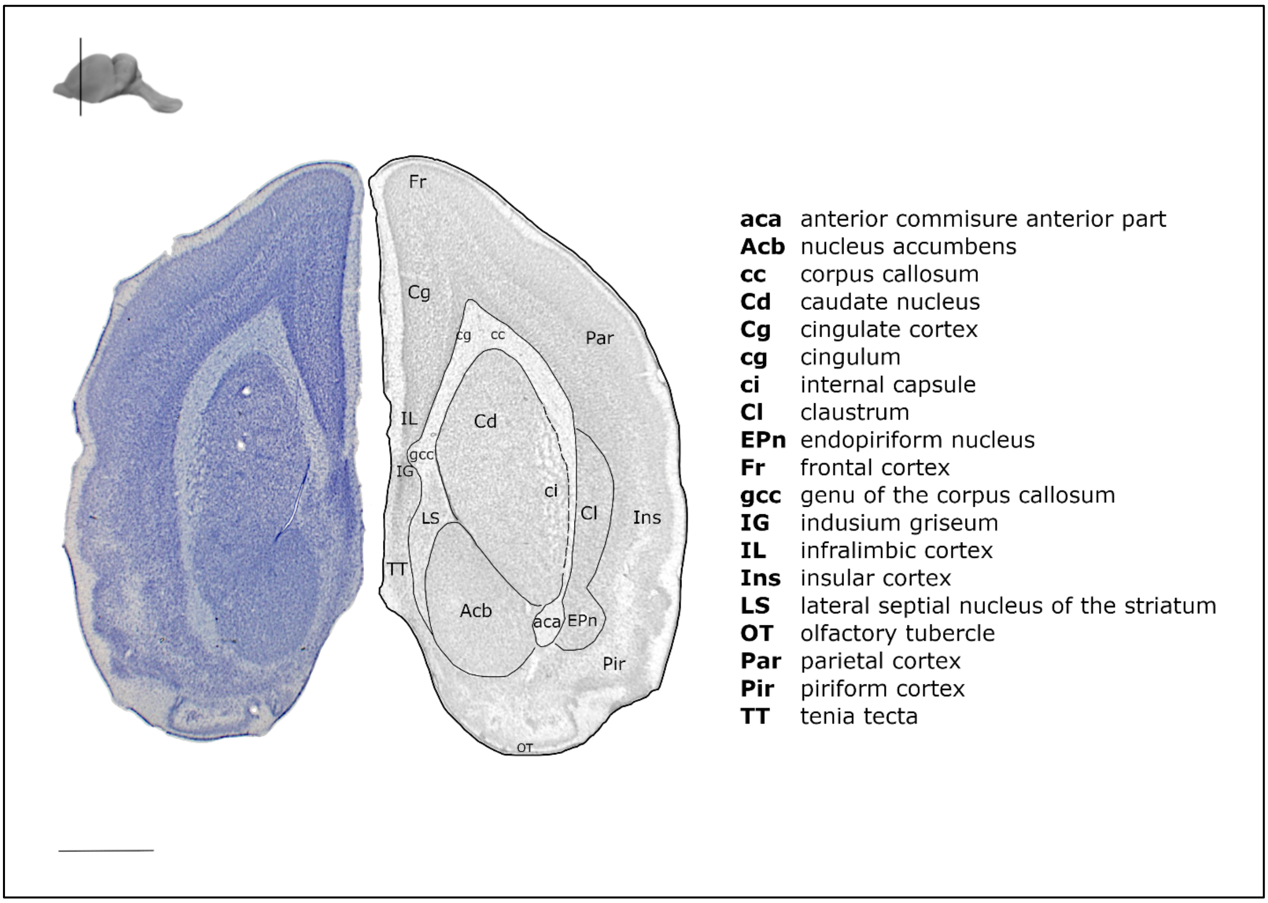

***Figure S 1 Nissl-stained brain coronal brain section of adult male CP individual A.*** *The section was derived from the anterior brain block generated by matrix guided blocking using the species-specific brain matrix followed by freezing microtoming. The brain blocks were generated using blade channel 3 of the sectioning matrix. For annotation brain atlases for the bat species CP (Scalia et al., 2013), Phyllostomus discolor (Radtke-Schuller et al., 2020), Rousettus aegyptiacus (Eilam-Altstadter et al., 2021), as well as the Alen mouse brain atlas (Allen Mouse Brain Atlas, mouse.brain-map.org and atlas.brain-map.org) were used as reference. Size bar = 1 mm.*

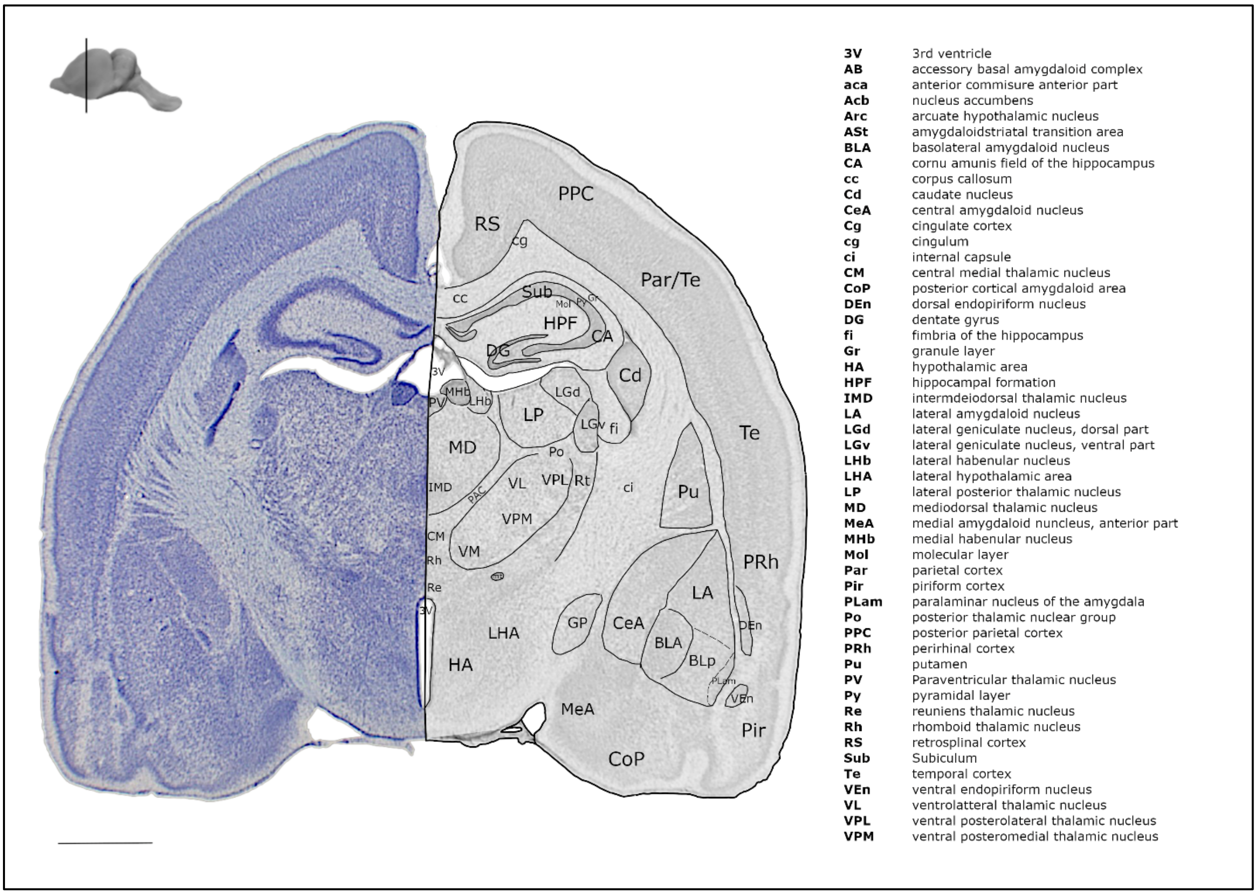

***Figure S 2 Nissl-stained brain coronal brain section of adult male CP individual A.*** *The section was derived from the anterior brain block generated by matrix guided blocking using the species-specific brain matrix followed by freezing microtoming. The brain blocks were generated using blade channel 3 of the sectioning matrix. For annotation brain atlases for the bat species CP (Scalia et al., 2013), Phyllostomus discolor (Radtke-Schuller et al., 2020), Rousettus aegyptiacus (Eilam-Altstadter et al., 2021), as well as the Alen mouse brain atlas (Allen Mouse Brain Atlas, mouse.brain-map.org and atlas.brain-map.org) were used as reference. Size bar = 1 mm.*

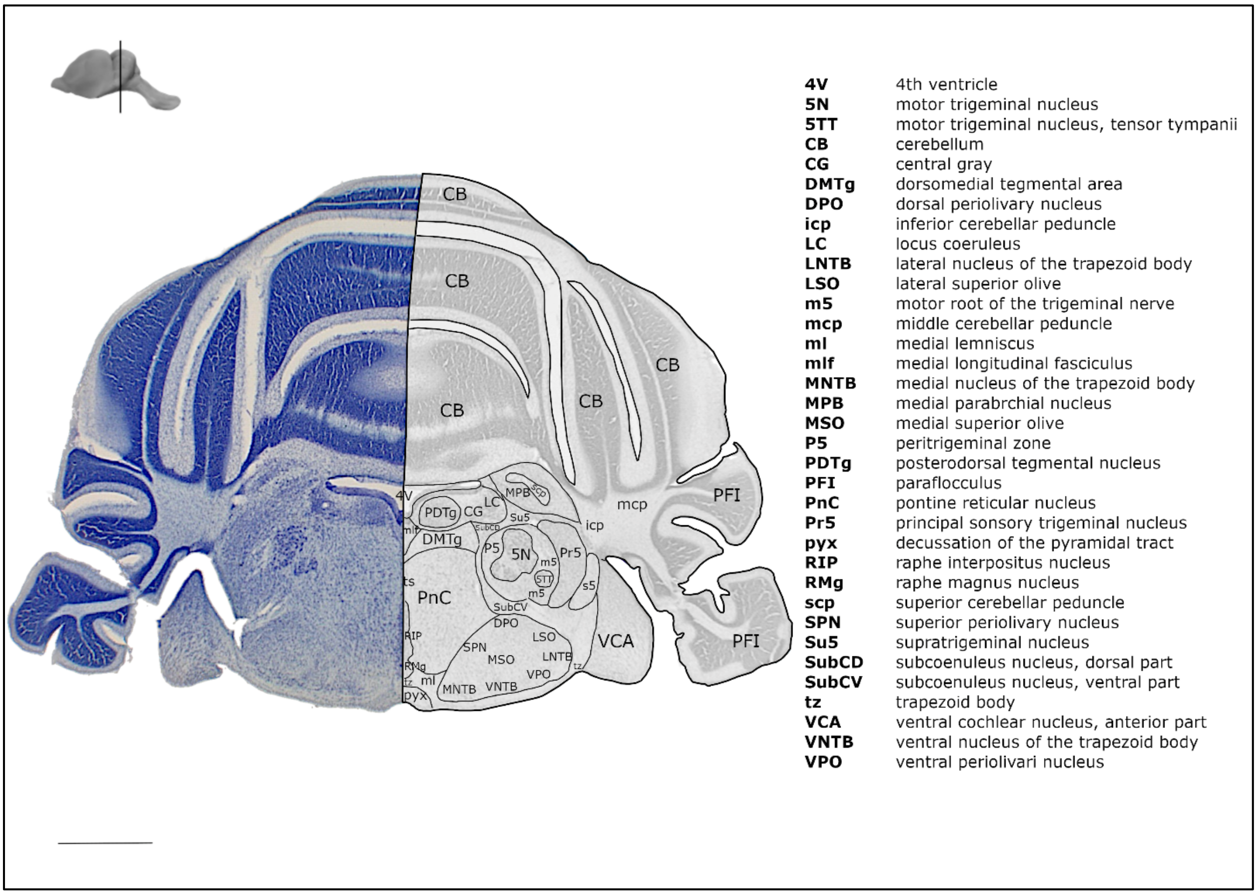

***Figure S 3 Nissl-stained brain coronal brain section of adult male CP individual A.*** *The section was derived from the posterior brain block generated by matrix guided blocking using the species-specific brain matrix followed by freezing microtoming. The brain blocks were generated using blade channel 3 of the sectioning matrix. For annotation brain atlases for the bat species CP (Scalia et al., 2013), Phyllostomus discolor (Radtke-Schuller et al., 2020), Rousettus aegyptiacus (Eilam-Altstadter et al., 2021), as well as the Alen mouse brain atlas (Allen Mouse Brain Atlas, mouse.brain-map.org and atlas.brain-map.org) were used as reference. Size bar = 1 mm.*

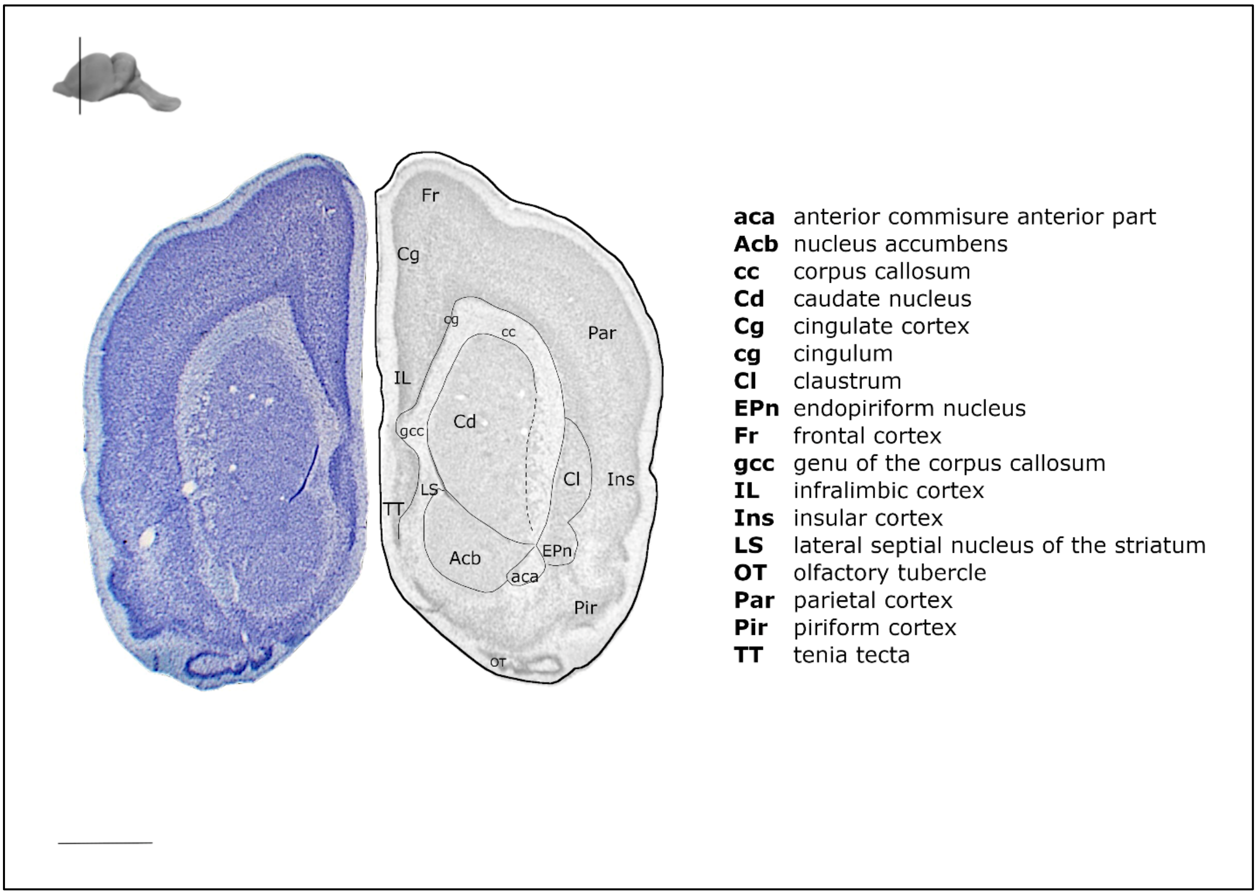

***Figure S 4 Nissl-stained brain coronal brain section of adult male CP individual B.*** *The section was derived from the anterior brain block generated by matrix guided blocking using the species-specific brain matrix followed by freezing microtoming. The brain blocks were generated using blade channel 3 of the sectioning matrix. For annotation brain atlases for the bat species CP (Scalia et al., 2013), Phyllostomus discolor (Radtke-Schuller et al., 2020), Rousettus aegyptiacus (Eilam-Altstadter et al., 2021), as well as the Alen mouse brain atlas (Allen Mouse Brain Atlas, mouse.brain-map.org and atlas.brain-map.org) were used as reference. Size bar = 1 mm.*

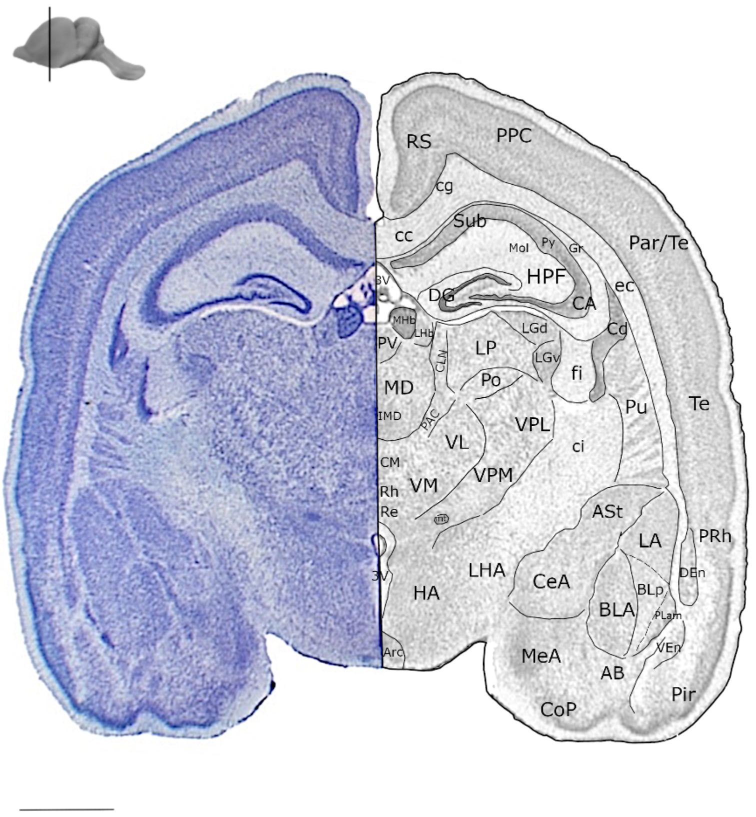

| | |
|---|---|
| **3V** | 3rd ventricle |
| **AB** | accessory basal amygdaloid complex |
| **aca** | anterior commisure anterior part |
| **Acb** | nucleus accumbens |
| **Arc** | arcuate hypothalamic nucleus |
| **ASt** | amygdaloidstriatal transition area |
| **BLA** | basolateral amygdaloid nucleus |
| **CA** | cornu amunis field of the hippocampus |
| **cc** | corpus callosum |
| **Cd** | caudate nucleus |
| **CeA** | central amygdaloid nucleus |
| **Cg** | cingulate cortex |
| **cg** | cingulum |
| **ci** | internal capsule |
| **CM** | central medial thalamic nucleus |
| **CoP** | posterior cortical amygdaloid area |
| **DEn** | dorsal endopiriform nucleus |
| **DG** | dentate gyrus |
| **fi** | fimbria of the hippocampus |
| **Gr** | granule layer |
| **HA** | hypothalamic area |
| **HPF** | hippocampal formation |
| **IMD** | intermdeiodorsal thalamic nucleus |
| **LA** | lateral amygdaloid nucleus |
| **LGd** | lateral geniculate nucleus, dorsal part |
| **LGv** | lateral geniculate nucleus, ventral part |
| **LHb** | lateral habenular nucleus |
| **LHA** | lateral hypothalamic area |
| **LP** | lateral posterior thalamic nucleus |
| **MD** | mediodorsal thalamic nucleus |
| **MeA** | medial amygdaloid nuncleus, anterior part |
| **MHb** | medial habenular nucleus |
| **Mol** | molecular layer |
| **Par** | parietal cortex |
| **Pir** | piriform cortex |
| **PLam** | paralaminar nucleus of the amygdala |
| **Po** | posterior thalamic nuclear group |
| **PPC** | posterior parietal cortex |
| **PRh** | perirhinal cortex |
| **Pu** | putamen |
| **PV** | Paraventricular thalamic nucleus |
| **Py** | pyramidal layer |
| **Re** | reuniens thalamic nucleus |
| **Rh** | rhomboid thalamic nucleus |
| **RS** | retrosplinal cortex |
| **Sub** | Subiculum |
| **Te** | temporal cortex |
| **VEn** | ventral endopiriform nucleus |
| **VL** | ventrolatteral thalamic nucleus |
| **VPL** | ventral posterolateral thalamic nucleus |
| **VPM** | ventral posteromedial thalamic nucleus |

***Figure S 5 Nissl-stained brain coronal brain section of adult male CP individual B.*** *The section was derived from the anterior brain block generated by matrix guided blocking using the species-specific brain matrix followed by freezing microtoming. The brain blocks were generated using blade channel 3 of the sectioning matrix. For annotation brain atlases for the bat species CP (Scalia et al., 2013), Phyllostomus discolor (Radtke-Schuller et al., 2020), Rousettus aegyptiacus (Eilam-Altstadter et al., 2021), as well as the Alen mouse brain atlas (Allen Mouse Brain Atlas, mouse.brain-map.org and atlas.brain-map.org) were used as reference. Size bar = 1 mm.*

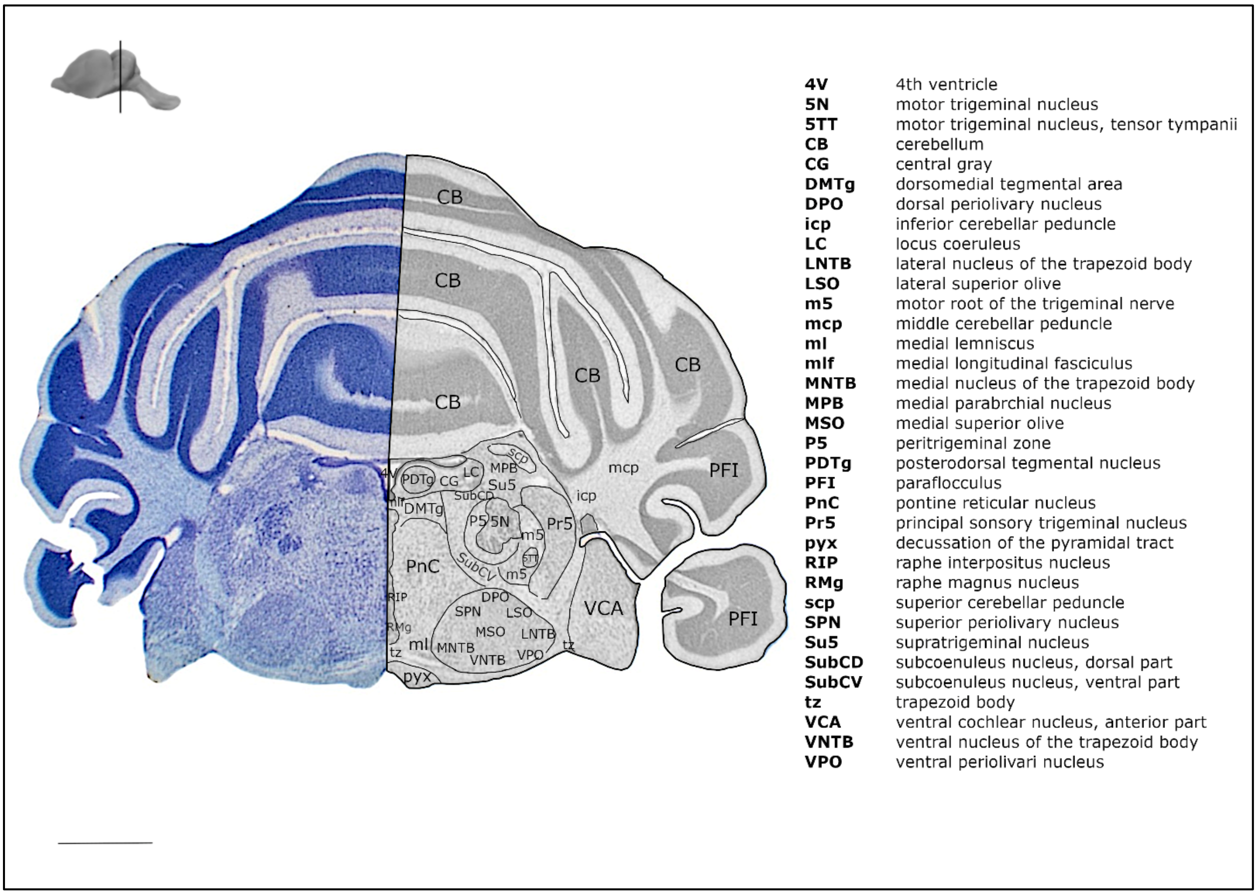

***Figure S 6 Nissl-stained brain coronal brain section of adult male CP individual B.*** *The section was derived from the posterior brain block generated by matrix guided blocking using the species-specific brain matrix followed by freezing microtoming. The brain blocks were generated using blade channel 3 of the sectioning matrix. For annotation brain atlases for the bat species CP (Scalia et al., 2013), Phyllostomus discolor (Radtke-Schuller et al., 2020), Rousettus aegyptiacus (Eilam-Altstadter et al., 2021), as well as the Alen mouse brain atlas (Allen Mouse Brain Atlas, mouse.brain-map.org and atlas.brain-map.org) were used as reference. Size bar = 1 mm.*
